# Sirt7 regulates circadian phase coherence of hepatic circadian clock via a body temperature/Hsp70-Sirt7-Cry1 axis

**DOI:** 10.1101/363176

**Authors:** Zuojun Liu, Minxian Qian, Xiaolong Tang, Wenjing Hu, Shuju Zhang, Fanbiao Meng, Shimin Sun, Xinyue Cao, Qiuxiang Pang, Bosheng Zhao, Baohua Liu

## Abstract

The biological clock is generated in the hypothalamic suprachiasmatic nucleus (SCN), which synchronizes peripheral oscillators to coordinate physiological and behavioral activities throughout the body. Disturbance of circadian phase coherence between the central and peripheral could disrupt rhythms and thus cause diseases and aging. Here, we identified hepatic Sirt7 as an early element responsive to light, which ensures the phase coherence in mouse liver. Loss of *Sirt7* leads to advanced liver circadian phase; restricted feeding in daytime entrains hepatic clock more rapidly in *Sirt7-/-* mice compared to wild-types. Molecularly, a light-driven body temperature (BT) oscillation induces rhythmic expression of Hsp70, which binds to and promotes the ubiquitination and proteasomal degradation of Sirt7. Sirt7 rhythmically deacetylates Cry1 on K565/579 and promotes Fbxl3-mediated degradation, thus coupling hepatic clock to the central pacemaker. Together, our data identify a novel BT/Hsp70-Sirt7-Cry1 axis, which transmits biological timing cues from the central to the peripheral and ensures circadian phase coherence in livers.

## Introduction

The physiological and behavioral activities throughout our body are coordinated by an endogenous biological clock, which is generated by a central pacemaker in the SCN (Bass and Takahashi, 2010; Panda, 2016). Molecularly, circadian rhythmicity is sustained by an auto-regulatory transcriptional/ translational feedback loop (TTFL), in which Clock and Bmal1 heterodimerize to promote transcription of *Cryptochrome* (*Cry1/2*) and *Period* (*Per1/2*); accumulating Crys and Pers dimerize and translocate to the nucleus to repress the transcriptional activity of Clock:Bmal1 complex (Bunger et al., 2000; DeBruyne et al., 2007; Gekakis et al., 1998; Kume et al., 1999; Reick et al., 2001; Takahashi et al., 2008; van der Horst et al., 1999). In addition, the Clock:Bmal1 complex activates transcription of *Rev*-*Erbα/β* and retinoic acid receptor-related orphan receptor (*Ror*); Ror then stimulates but Rev-Erbα/β suppresses *Bmal1* transcription via ROR elements (RORE) (Preitner et al., 2002; Sato et al., 2004; Solt et al., 2012). The endogenous clock is entrained by a variety of environmental signals, i.e. *Zeitgebers*. Light represents a predominant *Zeitgeber*, which entrains the central clock in the SCN via the retino-hypothalamic tract; SCN then synchronizes subsidiary oscillators in peripheral (Dibner et al., 2010a). Though molecularly not very clear yet, increasing evidence suggests that such synchronization is mediated by systemic cues like feeding activity, BT and hormones (Mohawk et al., 2012b). BT decreases in the light, resting phase, but increases in the dark, active phase in mice (Gerhart-Hines et al., 2013; Orozco-Solis et al., 2016), integrating the central clock to hepatic clock via Heat shock factor (Hsf1)-mediated stress response (Brown et al., 2002; Buhr et al., 2010; Saini et al., 2012). Of note, the light/dark (LD) cycle also regulates food intake behavior, coupling the central clock to metabolism (Damiola et al., 2000; Fonken et al., 2010; Stokkan et al., 2001); feeding serves as an independent *Zeitgeber* of hepatic clock but not the central clock (Dibner et al., 2010a; Mohawk et al., 2012b). At molecular level, NAD^+^-dependent sirtuins are critical nutrient sensors that couple metabolism to peripheral clocks. Sirt1 rhythmically binds to and deacetylates Bmal1 to enhance the transcription activity of the Clock:Bmal1 complex (Nakahata et al., 2008). It also deacetylates and thus to destabilize Per2, and modulates both the central and peripheral clocks (Asher et al., 2008). Interestingly, Sirt6 rather regulates the expression of clock-controlled genes (CCGs) instead of core TTFL elements in the liver (Masri et al., 2014). Another member Sirt7 localizes in the nucleolus, which deacetylates H3K18 and desuccinylates H3K122 to modulate chromatin remodeling, thus regulating gene transcription and DNA repair (Barber et al., 2012; Li et al., 2016). Loss of *Sirt7* causes genomic instability, fatty liver and accelerated aging in mice (Vazquez et al., 2016). The biological function of Sirt7 in circadian rhythms are still largely unclear.

All environment *Zeitgebers*, the central and peripheral oscillators have to be finely aligned to ensure the circadian phase coherence, otherwise it might lead to disrupted rhythmicity, metabolic diseases and aging, e.g. constant light exposure could alter the feeding activity and causes obesity (Fonken et al., 2010). Ablating *Clock* in mice disrupts feeding rhythms and metabolism and accelerates aging (Dubrovsky et al., 2010; Turek et al., 2005), and *Bmal1* deleted mice are arrhythmic and short-lived (Bunger et al., 2000; Kondratov et al., 2006). Intriguingly, only marginal effects were observed on the rhythmicity of *Pers* and *Crys* in livers wherein the expression of *Bmal1* was attenuated (Kornmann et al., 2007). Restricted feeding regimen entrains hepatic clock more rapidly in mice ablated SCN (Saini et al., 2013). These suggest that synchronization cues from the central pacemaker counteract phase entrainment of liver clock mediated by feeding, in a Bmal1-independent yet unknown mechanism. Here we found that Sirt7 is an early element responsive to light-derived timing cue in mouse liver. Body temperature (BT) oscillation induces rhythmic transcription of *Hsp70*, which interacts with Sirt7 to promote its ubiquitination and proteasomal degradation. The rhythmic Sirt7 then deacetylates the core circadian oscillator Cry1 and promotes its degradation mediated by Fbxl3, coupling hepatic clock to the central pacemaker. Collectively, our data reveal a novel BT/Hsp70-Sirt7-Cry1 axis, which transmits the timing cues from the central to periphery and ensures circadian phase coherence in livers.

## Results

### Light-entrained BT oscillation generates Sirt7 rhythmicity in mouse liver

NAD^+^-dependent sirtuins are essential nutrient sensors that bridge circadian rhythms and metabolism in peripheral tissues like liver (Houtkooper et al., 2012), while their nutrient-independent roles in circadian rhythms is limited to date. We first examined the oscillation pattern of sirtuins across a LD cycle in mouse liver, the largest organ sensitive to feeding rhythms and widely applied to the study of peripheral oscillators. As shown, the rhythmicity of Bmal1 and Cry1 was prominent, and similarly, Sirt1, 3, 5, 6 and 7 exhibited obvious oscillation (Fig. 1A,B). Of particular interest, the levels of Sirt7 and Cry1 were tightly correlated across a LD cycle. Sirt7 accumulated during daytime but dropped at night around *Zeitgeber* time (ZT) 12, whereas Cry1 exhibited a completely inversed pattern. Of note, the mRNA level of *Sirt7* was lack of obvious oscillation (Supplemental Fig. S1A), and Sirt7 mRNA or protein was hardly detected in the hypothalamus (Supplemental Fig. S1B,C).

**Figure 1.**
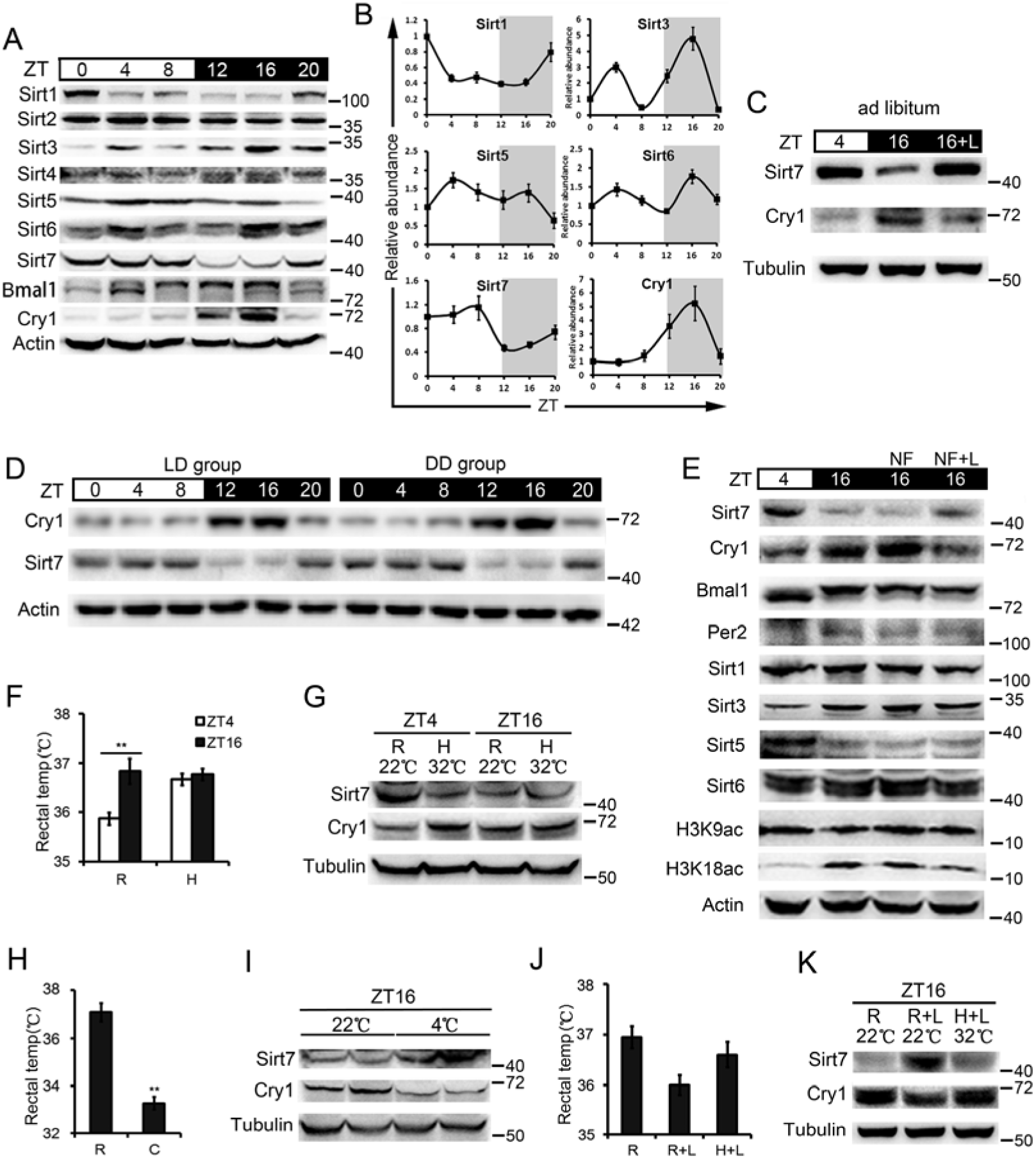
Light-entrained body temperature oscillation generates Sirt7 rhythmicity in mouse liver. (***A***) Western blots showing protein levels of Sirt1-7 and indicated core clock factors in wild-type (WT) mouse liver across a circadian period. Liver tissues were collected at indicated circadian times relative to Zeitgeber time (ZT). Representative blots from at least three independent experiments with different mice are shown. (***B***) Quantification of band intensity of three independent blots in (**A**) using Image J^®^. Data represent the means ± SEM. (***C***) Western blots showing Sirt7 and Cry1 protein levels under normal feeding condition with or without 2-h light (L) exposure from ZT14 to ZT16 in mouse livers. (***D***) Western blots showing levels of Sirt7 and Cry1 in 12:12 LD or dark/dark (DD) cycle in mouse livers. DD group was placed in darkness for 24 h, and liver tissues were collected every 4 h for 24 h. (n = 3 mice/genotype/time point). (***E***) Western blots showing indicated protein levels under normal feeding or fasting (no food, NF) conditions with or without 2-h light exposure (L). Mice were fasted from ZT0 to ZT16 under a normal light/dark (LD) cycle or exposed to a 2-h light pulse from ZT14 to ZT16 (n = 3 mice/genotype/time point). (***F***) Mouse rectal temperature under room (R) or high (H) ambient temperature at ZT4 and ZT16. Data represent the means ± SEM of three independent experiments. \*\**P* < 0.01. (***G***) Western blots showing Sirt7 and Cry1 protein levels under room (R) or high (H) ambient temperatures at ZT4 and ZT16 (n = 3 mice/genotype/time point). (***H***) Mouse rectal temperature under room or cold (C) ambient temperature at ZT16. Data represent the means ± SEM of three independent experiments. \*\**P* < 0.01. (***I***) Western blots showing protein levels of Sirt7 and Cry1 at ZT16 under room or cold ambient temperature at ZT16 (n = 3 mice/genotype/time point). (***J***) Mouse rectal temperature with 2-h light exposure under R or H ambient temperature at ZT16 (n = 3 mice/genotype/time point). Data represent the means ± SEM of three independent experiments. \*\**P* < 0.01. (***K***) Western blots showing protein levels of Sirt7 and Cry1 in mouse liver with 2-h light exposure under room or high ambient temperature at ZT16 (n = 3 mice/genotype/time point).

Light represents the most prominent *Zeitgeber* of circadian clock. To further decipher the effect of LD, we examined the hepatic level Sirt7 at ZT4 (day) and ZT16 (night) with or without a 2-h light exposure from ZT14 to ZT16 (L) in dark period in mice fed *ad libitum*. As shown, a dramatic reduction in Sirt7 level was observed at ZT16 compared to ZT4. Moreover, light exposure in the dark elevated the level of Sirt7 but decreased that of Cry1 (Fig. 1C), suggesting that LD modulates Sirt7 level in the liver. To examine whether Sirt7 was clock-controlled rather than induced by physiological changes upon light exposure, we switched the mice from LD cycle to dark/dark (DD) cycle. Indeed, consistent with the self-sustained property of circadian clock, levels of Sirt7 and Cry1 were oscillated in a circadian manner in DD, similar to that in LD (Fig. 1D). The data support the notion that Sirt7 belongs to clock-controlled genes.

LD regulates feeding activity (Fonken et al., 2010), which entrains hepatic clock but hardly affects the central pacemaker (Dibner et al., 2010b). Given the lack of Sirt7 in the hypothalamus, we hypothesized that light regulates hepatic Sirt7 via systemic cues like feeding-fasting cycle (FF). To test the possibility, we examined Sirt7 level at ZT4 and ZT16 under feeding or fasting condition (no feeding from ZT0 to ZT16, NF), with a normal LD cycle or a 2-h light exposure (ZT14 to ZT16). While fasting hardly affected the level of hepatic Sirt7 in the dark period (ZT16NF Vs ZT16), the 2-h light exposure remarkably increased Sirt7 level (ZT16NF+L Vs ZT16NF) (Fig. 1E). This suggests that light resets the expressing pattern of Sirt7 independent of feeding. In line with the rhythmic expression of Sirt7, the level of H3K18ac, a direct target of Sirt7 deacetylase (Barber et al., 2012), exhibited a totally inversed oscillating pattern. By contrast, the levels of Sirt1, Sirt6 and their deacetylating target H3K9ac between ZT4 and ZT16 were merely changed, regardless of feeding, fasting or light exposure.

Though oscillation of Sirt3, Sirt5, and Bmal1 was obvious between ZT4 and ZT16 in mice fed *ad libitum*, minimal effect was observed under light and/or fasting condition. Of note, mRNA levels of *Sirt7*, *Cry1*, *Bmal1* and *Per2* in the liver were merely affected by light exposure in the dark period (Supplemental Fig. S1D). Collectively, the data indicate that the rhythmicity of hepatic Sirt7 is entrained in a FF-independent manner.

In addition to FF cycle, body temperature (BT) is also a systemic cue employed by the SCN to synchronize peripheral clocks (Buhr et al., 2010; Mohawk et al., 2012b). BT can be elevated at environmental temperatures exceeded the thermoneutral zone (30°C) in mice (Fischer et al., 2016). Indeed, alteration of ambient temperature (AT) imparts changes on circadian gene expression in peripheral organs (Brown et al., 2002). We found that BT oscillation was maintained in DD (Supplemental Fig. S2A). We thus asked whether Sirt7 oscillation is regulated by BT. We subjected the mice to a high AT (32°C) at different phases of a circadian cycle and measured rectal temperature (RT). Under room temperature, RT was high at ZT16 (36.9 ± 0.3°C) and low at ZT4 (35.9 ± 0.1°C) (Fig. 1F). High AT immediately enhanced RT (36.7 ± 0.1°C Vs 35.9 ± 0.1°C) at ZT4 but not at ZT16, and significantly downregulated the level of Sirt7 but increased that of Cry1 at ZT4 (Fig. 1G). By contrast, cold treatment (4°C) at ZT16 decreased RT (37.1 ± 0.3°C Vs 33.3 ± 0.3°C), increased Sirt7, and downregulated Cry1 (Fig. 1H,I). Next, we examined whether light exposure at night stabilizes Sirt7 via BT. As shown, a 2-h light exposure decreased RT, and that was blocked by high AT challenge (Fig. 1J). Importantly, light-induced increase of Sirt7 level at ZT16 was abolished by a concomitant high AT challenge (Fig. 1K). Given that BT rhythmicity can be indirectly regulated by feeding and locomotor activity (Mohawk et al., 2012a), we further examined the effect of fasting on BT oscillation. Indeed, the BT slightly declined at fasting condition, but further dramatically decreased upon light exposure at ZT16 in fasted mice (Supplemental Fig. 2C,D), supporting the notion that the Sirt7 oscillation was regulated independent of FF cycle. Together, these results suggest that light-entrained BT oscillation underlies rhythmic expression of hepatic Sirt7 independent of feeding.

**Figure 2.**
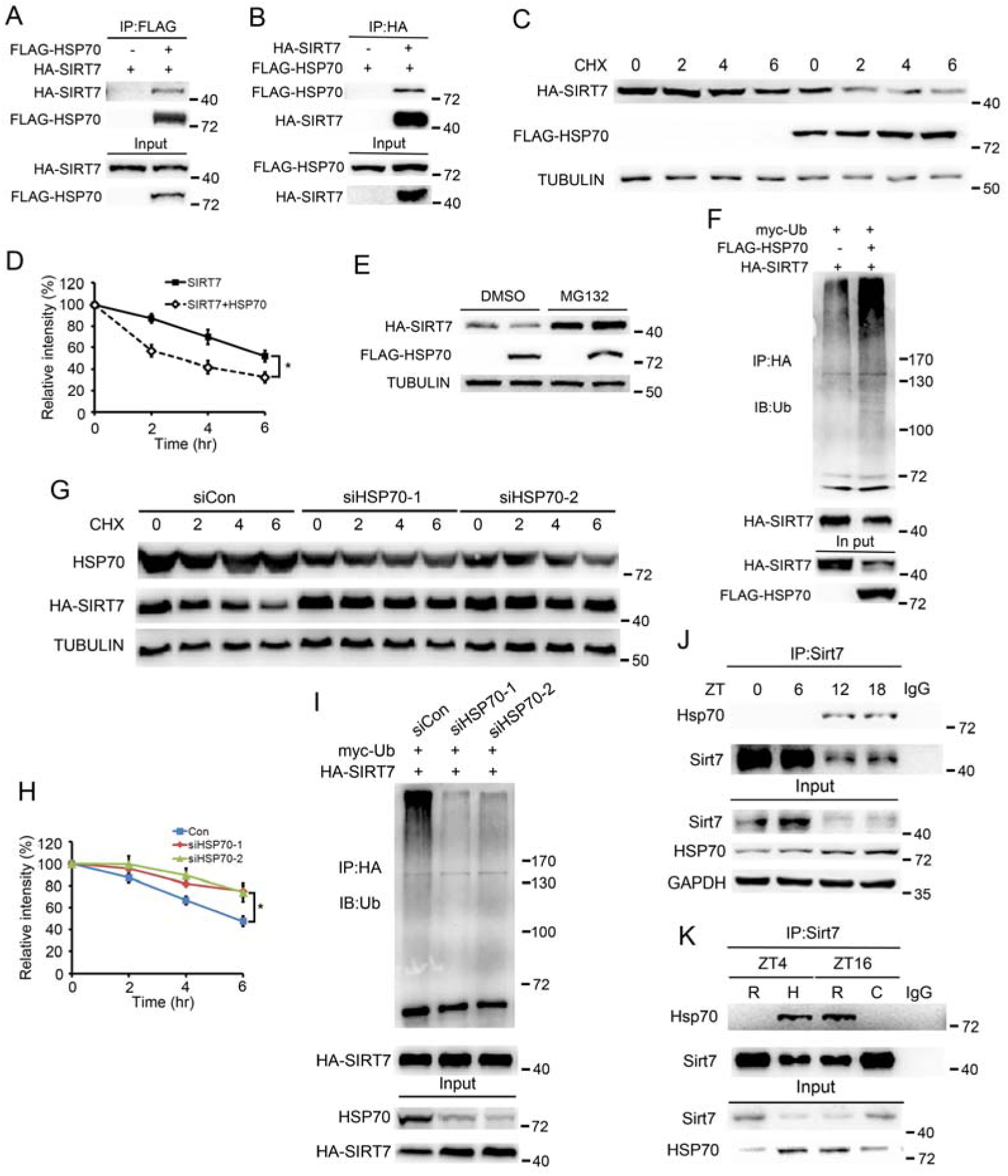
Hsp70 mediates Sirt7 degradation via ubiquitin-proteasome pathway. (***A*-*B***) Co-immunoprecipitation (IP) was performed in HEK293 cells over-expressing HA-Sirt7 and FLAG-Hsp70. (***C***) Western blots showing Sirt7 levels in HEK293 cells expressing FLAG-Hsp70 and HA-Sirt7 treated with 50 mg/ml cycloheximide (CHX). (***D***) Quantification of Sirt7 levels in (**C**) by Image J^®^. Data represent the means ± SEM of three independent experiments. \**P* < 0.05. (***E***) Representative Western blots showing Sirt7 levels in HEK293 cells expressing FLAG-Hsp70 and HA-Sirt7 treated with 20 μM MG132 for 6 h. (***F***) Ubiquitination of Sirt7 in the absence or presence of FLAG-Hsp70. (***G***) Western blots showing Sirt7 levels in Scramble (siCon) or siHsp70 treated HEK293 cells in the presence of CHX (50□μg/ml). (***H***) Quantification of Sirt7 levels in (**G**). Data represent the means ± SEM of three independent experiments. \**P* < 0.05. (***I***) Ubiquitination of Sirt7 in Scramble or siHsp70-treated HEK293 cells. (***J***) The interaction between Hsp70 and Sirt7 in mouse livers across a circadian cycle was examined by Co-IP. (***K***) The interaction between Hsp70 and Sirt7 at various ATs was determined by Co-IP.

### A Hsp70-Sirt7 axis transmits synchronization cues to mouse liver

BT resets peripheral clocks via transcription factor Hsf1 and downstream heat-shock proteins (Hsps), i.e. Hsp70, Hspca, Hsp105 and Hspa8 etc. (Kornmann et al., 2007; Saini et al., 2012). We found that the transcription of *Hsps* was significantly enhanced by high AT challenge at ZT4 (Supplemental Fig. S2B). Hsp70 is a key chaperone that regulates protein homeostasis (Balchin et al., 2016). Considering that BT-driven Sirt7 oscillation (see Fig. 1, high at ZT4 Vs low at ZT16) is negatively correlated with Hsp70 level (Supplemental Fig. S2, low at ZT4 Vs high at ZT16), we asked whether BT regulates Sirt7 rhythmicity via Hsp70. To test it, co-immunoprecipitation (Co-IP) was first applied to determine potential interaction between Sirt7 and Hsp70 proteins. As shown, Sirt7 interacted with Hsp70 (Fig. 2A,B). Further, degradation assay revealed that overexpression of Hsp70 accelerated the degradation of HA-Sirt7 (Fig. 2C,D), which was blocked by proteasome inhibitor MG132 (Fig. 2E). Hsp70 overexpression increased the poly-ubiquitination of HA-Sirt7 (Fig. 2F). Knocking down (KD) *Hsp70* via siRNA suppressed the degradation of HA-Sirt7 (Fig. 2G,H) and downregulated poly-ubiquitination of HA-Sirt7 (Fig. 2I). These data indicate that Hsp70 promotes Sirt7 degradation via ubiquitin-proteasome system.

To verify the *in vivo* function of Hsp70, endogenous Sirt7 was immunoprecipitated from the liver lysates across a circadian cycle. A rhythmic interaction between Hsp70 and Sirt7 was observed with maximum binding at ZT12 and ZT16, consistent with high level of Hsp70 but low level of Sirt7 in the dark period (Fig. 2J). Next, we asked whether the interaction between Hsp70 and Sirt7 was modulated by BT. To this end, the interaction between Hsp70 and Sirt7 at different AT was examined. Interestingly, while high AT increased the binding of Hsp70 to Sirt7 at ZT4, low AT reduced such interaction at ZT16 (Fig. 2K).

We then examined whether Hsp70 is required for BT-generated Sirt7 rhythmicity. We first investigated whether heat shock inhibits Sirt7 expression at cellular level. As shown, a 2-h heat shock (39.5°C) led to reduced protein level of Sirt7 but increased Hsp70 and Cry1 in mouse embryonic fibroblasts (MEF) (Fig. 3A). Then, we knocked down *Hsp70* via siRNA. While the protein level of Sirt7 was elevated, that of Cry1 was decreased in *Hsp70* KD MEFs; moreover, the effects of heat shock on Sirt7 and Cry1 were blocked by *Hsp70* KD, suggesting that temperature relies on Hsp70 to regulate Sirt7 protein stability. We further investigated whether temperature regulates Cry1 via Sirt7. To this end, we generated *Sirt7* KO mice via the CRISPR/Cas9 procedure. Intriguingly, loss of *Sirt7* significantly upregulated Cry1 in MEFs (Fig. 3B). While the 2-h heat shock induced Hsp70 expression in both wild-type (WT) and *Sirt7*-/- cells, the increase of Cry1 was only observed in WT. We next applied various temperature challenges to *Sirt7*-/- and WT mice. Again, an upregulation of Cry1 was observed in *Sirt7*-/- livers (Fig. 3C,E). Consistent with the *in vitro* data, depletion of *Sirt7* completely abolished high AT challenge-induced upregulation of Cry1 at ZT4 in mouse livers (Fig. 3C,D). Similarly, cold treatment-induced Cry1 downregulation was totally blocked in *Sirt7*-/- livers (Fig. 3E,F). In addition, the expression of Hsp70 was comparable in WT and *Sirt7* KO mice liver at ZT16 by light exposure (Supplemental Fig, 2E,F). Together, the data support an Hsp70-Sirt7 axis, which transmits systemic synchronization cues to liver clock.

**Figure 3.**
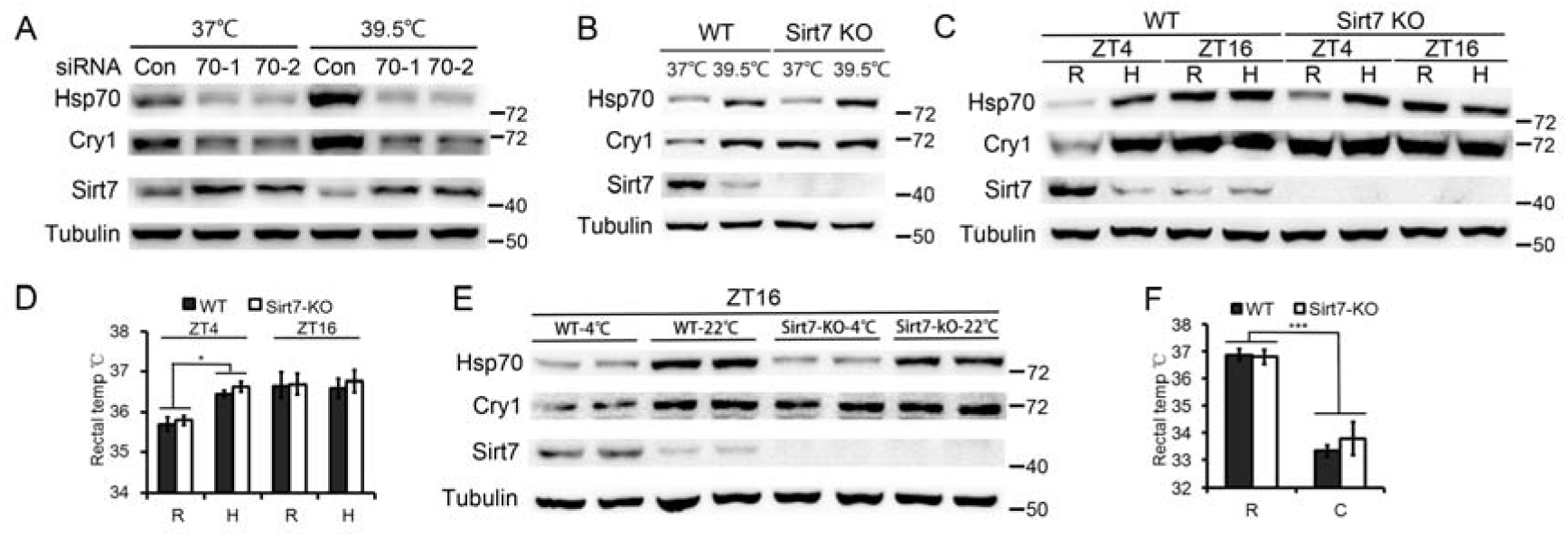
BT regulates Sirt7 level via Hsp70. (***A***) Western blots showing Sirt7, Hsp70 and Cry1 protein levels in WT and *Hsp70* KD MEFs with or without 2-h heat shock at 39.5 °C. (***B***) Western blots showing Sirt7, Hsp70 and Cry1 protein levels in WT and *Sirt7* KO MEFs with or without 2-h heat shock at 39.5 °C. (***C***) Western blots showing Sirt7, Hsp70 and Cry1 protein levels in WT and *Sirt7* KO liver tissues under room (R) or high (H) ambient temperatures at ZT4 and ZT16 (n = 3 mice/genotype/time point). (***D***) Mouse rectal temperature under room (R) or high (H) ambient temperature at ZT4 and ZT16. Data represent the means ± SEM of three independent experiments. \**P* < 0.05. (***E***) Western blots showing Sirt7, Hsp70 and Cry1 protein levels in WT and *Sirt7* KO liver tissues at ZT16 under room or cold ambient temperature at ZT16 (n = 3 mice/genotype/time point). (***F***) Mouse rectal temperature at ZT16 under room or cold ambient temperature at ZT16 (n = 3 mice/genotype/time point). Data represent the means ± SEM of three independent experiments. \*\*\**P* < 0.001.

### Sirt7 deacetylates Cry1

We repeatedly observed that the protein level of Cry1 was inversely correlated with that of Sirt7, peaking in the night but dropping in the day, and *Sirt7* deficiency led to a dramatic increase in Cry1 level in day time. We speculated that Sirt7 might regulate Cry1. We first examined whether Sirt7 and Cry1 interact with each other. To this end, HA-Sirt7 and FLAG-Cry1 were co-overexpressed in HEK293 cells. Co-IP experiment revealed that Sirt7 interacted with Cry1 (Fig. 4A,B). Similarly, endogenous Sirt7 was found in anti Cry1 immunoprecipitates (Fig. 4C), and the result of GST pull down assay supports a direct interaction between His-Cry1 and GST-Sirt7 (Fig. 4D).

**Figure 4.**
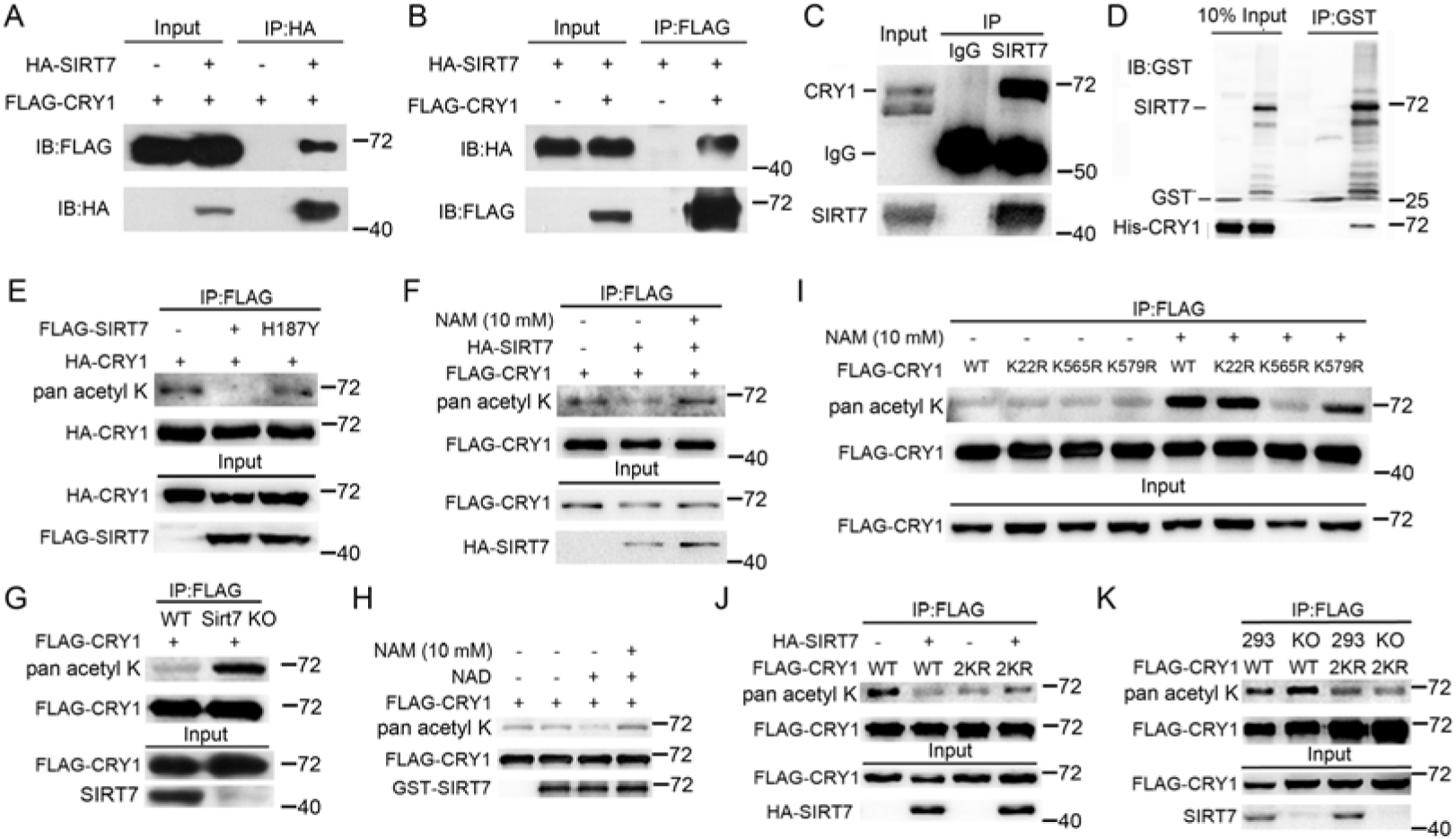
Sirt7 interacts with and deacetylates Cry1. (***A*-*B***) Co-immunoprecipitation (IP) was performed in HEK293 cells over-expressing HA-Sirt7 and FLAG-Cry1. (***C***) The interaction between endogenous Sirt7 and Cry1 was determined by Co-IP. (***D***) GST pull-down assay showing direct interaction between Sirt7 and Cry1. (***E***) Acetylation level of Cry1 in the presence of Sirt7 or catalytically inactive Sirt7 H187Y mutant. (***F***) Acetylation level of Cry1 in the presence of Sirt7 and/or 10 mM nicotinamide (NAM). (***G***) Western blots showing acetylation of FLAG-Cry1 in *SIRT7* KO HEK293 cells. (***H***) FLAG-Cry1 was immunoprecipitated and then incubated with purified GST-Sirt7, 1 mM NAD^+^ or 10 mM NAM. Acetylation of FLAG-Cry1 was detected with pan-acetyl K antibodies. (***I***) Western blots analysis of acetylation levels of WT and FLAG-Cry1 KR in the absence or presence of 10 mM NAM. KR mutant constructs have had lysine (K) residues mutated to an arginine (R) by site-directed mutagenesis. (***J***) Acetylation level of FLAG-Cry1-2KR in HEK293 cells overexpressing HA-Sirt7. (***K*)** Western blots showing acetylation levels of FLAG-Cry1-2KR in *Sirt7* KO cells.

As protein deacetylase (Kiran et al., 2015), Sirt7 might directly deacetylate Cry1. To test the hypothesis, we immunoprecipitated HA-Cry1 and performed Western blotting with pan acetyl K antibodies. As shown, the acetylation level of HA-Cry1 was remarkably reduced in the presence of Sirt7 but was unaffected in the presence of a catalytically inactive Sirt7 H187Y (Fig. 4E). The deacetylation of FLAG-Cry1 was abolished in cells treated with nicotinamide (NAM), a pan sirtuin inhibitor (Fig. 4F). Therefore, Sirt7 deacetylase activity is required for Cry1 deacetylation. Further, we found that the acetylation level of Cry1 was increased in *Sirt7* KO HEK293 cells (Fig. 4G). To confirm that Sirt7 directly targets Cry1, we performed an *in vitro* deacetylation assay with recombinant FLAG-Cry1 and GST-Sirt7 proteins. As shown, the acetylation level of FLAG-Cry1 was decreased in the presence of GST-Sirt7 and NAD^+^, which was blocked in the presence of NAM (Fig. 4H). Of note, although Cry2 is a close homologue of Cry1, the acetylation level of Cry2 was merely affected by NAM (Supplemental Fig. S3A). We also examined whether other sirtuins affected acetylation of Cry1. As shown, only Sirt7 strongly inhibited the acetylation of Cry1 (Supplemental Fig. S3B). These data suggest that Sirt7 specifically deacetylates Cry1.

To identify specific residues targeted for Sirt7-mediated deacetylation, we purified FLAG-Cry1 from HEK293 cells with or without ectopic HA-Sirt7 and did mass spectrometry. Three Ks were found differentially acetylated, i.e. K22/565/579 (Supplemental Fig. S4). We then generated nonacetylatable Cry1 mutants, wherein K22, K565 or K579 was replaced with Arginine (R). Each Cry1 mutant (FLAG-Cry1-KR) exhibited a comparable baseline acetylation level compared to WT (Fig. 4I). We then examined the effect of NAM on various FLAG-Cry1-KR mutants. As shown, NAM treatment enhanced the acetylation level of WT and K22R but had a minimal effect on K565R and only a moderate increase of K579R (Fig. 4I), indicating that K565/579 are targeted by Sirt7 deacetylation. We then generated FLAG-Cry1-2KR, wherein both K565 and K579 were mutated to Rs. As shown, FLAG-Cry1-2KR displayed much less acetylation than WT, and ectopic Sirt7 was unable to further downregulate the acetylation level (Fig. 4J). In addition, the acetylation level of FLAG-Cry1-2KR did not increase in *Sirt7* KO cells (Fig. 4K) or cells treated with NAM (Supplemental Fig. S5), indicating that Sirt7 deacetylates Cry1 precisely at K565 and K579.

### Sirt7-mediated deacetylation destabilizes Cry1

In unsynchronized *SIRT7* KO cells, *Cry1* mRNA level was slightly decreased but the protein level was largely increased (Supplemental Fig. S6A,B). By contrast, Cry1 protein level was decreased in *SIRT7*-overexpressing cells (Supplemental Fig. S6C). These indicate that SIRT7 most likely regulates Cry1 protein stability. We then examined the degradation rate of FLAG-Cry1 in HEK293 cells overexpressing HA-Sirt7 and treated with cycloheximide (CHX). The degradation rate of FLAG-Cry1 was accelerated in the presence of HA-Sirt7 compared to control (Fig. 5A,B), and Cry1 degradation was abolished in case of the FLAG-Cry1-2KR mutant (Fig. 5C,D). Similar phenomena were observed for endogenous Cry1 (Supplemental Fig. S7A-D). Moreover, the degradation rate of Cry1 was not affected by catalytically inactive mutant Sirt7 H187Y (Supplemental Fig. S7E). As such, we propose that Sirt7 regulates Cry1 protein stability.

**Figure 5.**
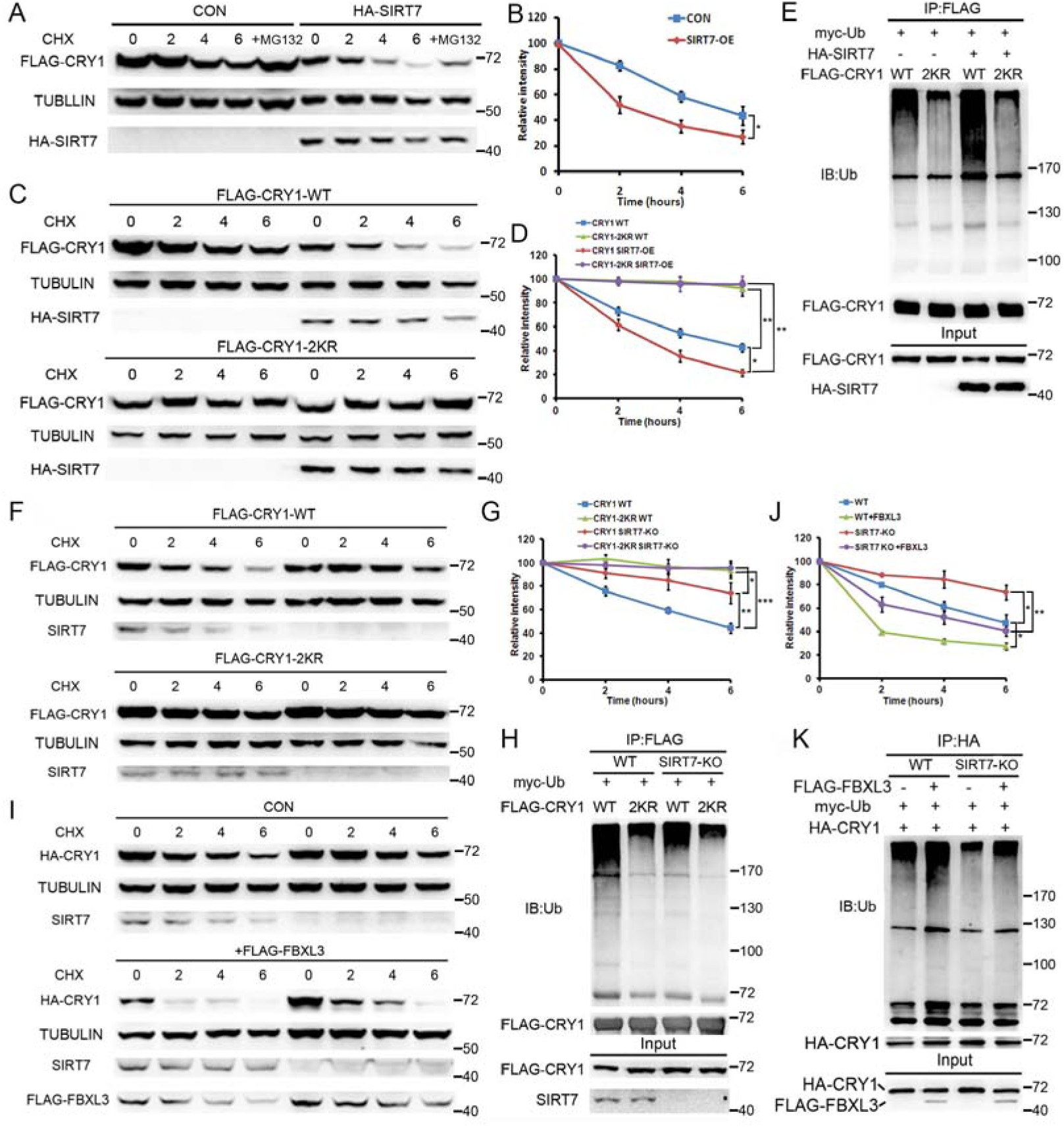
Sirt7-mediated deacetylation destabilizes Cry1. (***A***) Western blots showing Cry1 level in HEK293 cells expressing FLAG-Cry1 alone (control, CON) or co-overexpressing FLAG-Cry1 and HA-Sirt7, and treated with 50 mg/ml cycloheximide (CHX) and/or 20 μM MG132. (***B***) Quantification of Cry1 degradation in (**A**). Data represent the means ± SEM of three independent experiments. (***C***) Western blots showing degradation of FLAG-Cry1 and FLAG-Cry1-2KR in the presence or absence of HA-Sirt7. KR mutant constructs have had lysine (K) residues mutated to an arginine (R) by site-directed mutagenesis. (***D***) Quantification of Cry1 degradation in (**C**). Data represent the means ± SEM of three independent experiments. (***E***) Ubiquitination of Cry1 and Cry1-2KR in the absence or presence of HA-Sirt7. (***F***) Western blots showing the degradation rate of FLAG-Cry1 and FLAG-Cry1-2KR in *Sirt7* KO HEK293 cells generated by CRISPR/Cas9 procedure. (***G***) Quantification of Cry1 degradation in (**F**). Data represent the means ± SEM of three independent experiments. (***H***) Western blots showing ubiquitination (Ub) of FLAG-Cry1 and FLAG-Cry1-2KR in *Sirt7* KO HEK293 cells. (***I***) Western blots showing the effect of FBXL3 on Cry1 degradation in *Sirt7* KO HEK293 cells. (***J***) Quantification of Cry1 levels in (**I**). Data represent the means ± SEM of three independent experiments. (***K***) Western blots showing FBXL3-mediated ubiquitination of FLAG-Cry1 in *Sirt7*-KO HEK293 cells.

As Cry1 degradation was blocked by MG132, we reasoned that Sirt7 regulates Cry1 degradation via ubiquitination-proteasome pathway. Indeed, ectopic Sirt7 promoted poly-ubiquitination of Cry1 (Fig. 5E). Lysine acetylation increased protein stability by blocking ubiquitination on the same residue. K565/579 were previously identified as ubiquitination targets of F-box and leucine rich repeat protein 3 (Fbxl3), a subunit of ubiquitin protein ligase complex (SKP1-cullin-F-box, SCF) (Godinho et al., 2007; Hirano et al., 2013; Yoo et al., 2013). We speculated that Sirt7 deacetylates K565/579 to facilitate ubiquitination and subsequent degradation. As expected, ubiquitination level of FLAG-Cry1-2KR was dramatically reduced compared to WT and remained unchanged upon Sirt7 overexpression (Fig. 5E).

To determine whether acetylation stabilizes Cry1, we investigated the ubiquitination and degradation rate of Cry1 in *Sirt7*-/- cells. The degradation of FLAG-Cry1 was significantly suppressed in *Sirt7*-/- cells compared to control (Fig. 5F,G). By contrast, the degradation rate of FLAG-Cry1-2KR was comparable between *Sirt7*-/- and control cells. Moreover, the poly-ubiquitination levels of Cry1 were reduced in *Sirt7* KO cells (Fig. 5H). We asked whether increased acetylation protect Cry1 from Fbxl3-mediated degradation in *Sirt7* KO cells. We co-expressed FLAG-Fbxl3 and HA-Cry1 and analyzed the degradation rate. As expected, FLAG-Fbxl3 accelerated degradation of HA-Cry1, but this effect was significantly attenuated in *Sirt7* KO cells (Fig. 5I,J). We also examined Fbxl3-mediated poly-ubiquitination of Cry1 in *Sirt7* KO cells. Although the Cry1 poly-ubiquitination level was slightly increased upon Fbxl3 overexpression in *Sirt7* KO cells (which is likely attributable to other lysine residues), the overall ubiquitination level of Cry1 was significantly lower in *Sirt7* KO cells than controls (Fig. 5K). These results demonstrate that increased Cry1 protein stability in *Sirt7*-/- cells is due to elevated K565/579 acetylation, which prevents Fbxl3-mediated ubiquitination and subsequent proteasomal degradation. As AMPK regulates hepatic clock via phosphorylation-induced Cry1 degradation (Lamia et al., 2009), we thus examined whether K565/579 deacetylation affects AMPK-mediated Cry1 degradation. As shown (Supplemental Fig. S8), glucose starvation activated AMPK and it led to reduced protein levels of FLAG-Cry1 and FLAG-Cry1-2KR to a similar extent, indicating independent functions of Sirt7 and AMPK in regulating Cry1 stability.

### Sirt7 counteracts with circadian phase entrainment by feeding

The data suggests an essential role of Sirt7 in hepatic clocks. Deletion of *Sirt7* led to a dramatic increase of Cry1 at day time in mouse livers (Fig. 6A,B). We confirmed that hepatic Cry1 was acetylated in a cyclic manner, maximizing at ZT12-ZT18 in WT mice, but that was disrupted in *Sirt7*-/- mice showing elevated acetylation at ZT0-ZT6 (Fig. 6C,D). We therefore asked whether Sirt7 regulates circadian rhythms in a cell autonomous manner. As shown, compared to WT, reduced amplitude of *Bmal1*, *Cry1*, *Dbp*, *Rev*-*Erbα* and *Reverbβ* was observed in *Sirt7*-/- MEFs (Supplemental Fig. S9A). The protein level of Cry1 was elevated, whereas that of Bmal1 was reduced in *Sirt7*-/- cells (Supplemental Fig. S9B,C). Very interestingly, we found that circadian phase of *Per2* was significantly delayed in *Sirt7*-/- MEFs. Together, these data implicate that Sirt7 regulates endogenous circadian clocks.

**Figure 6.**
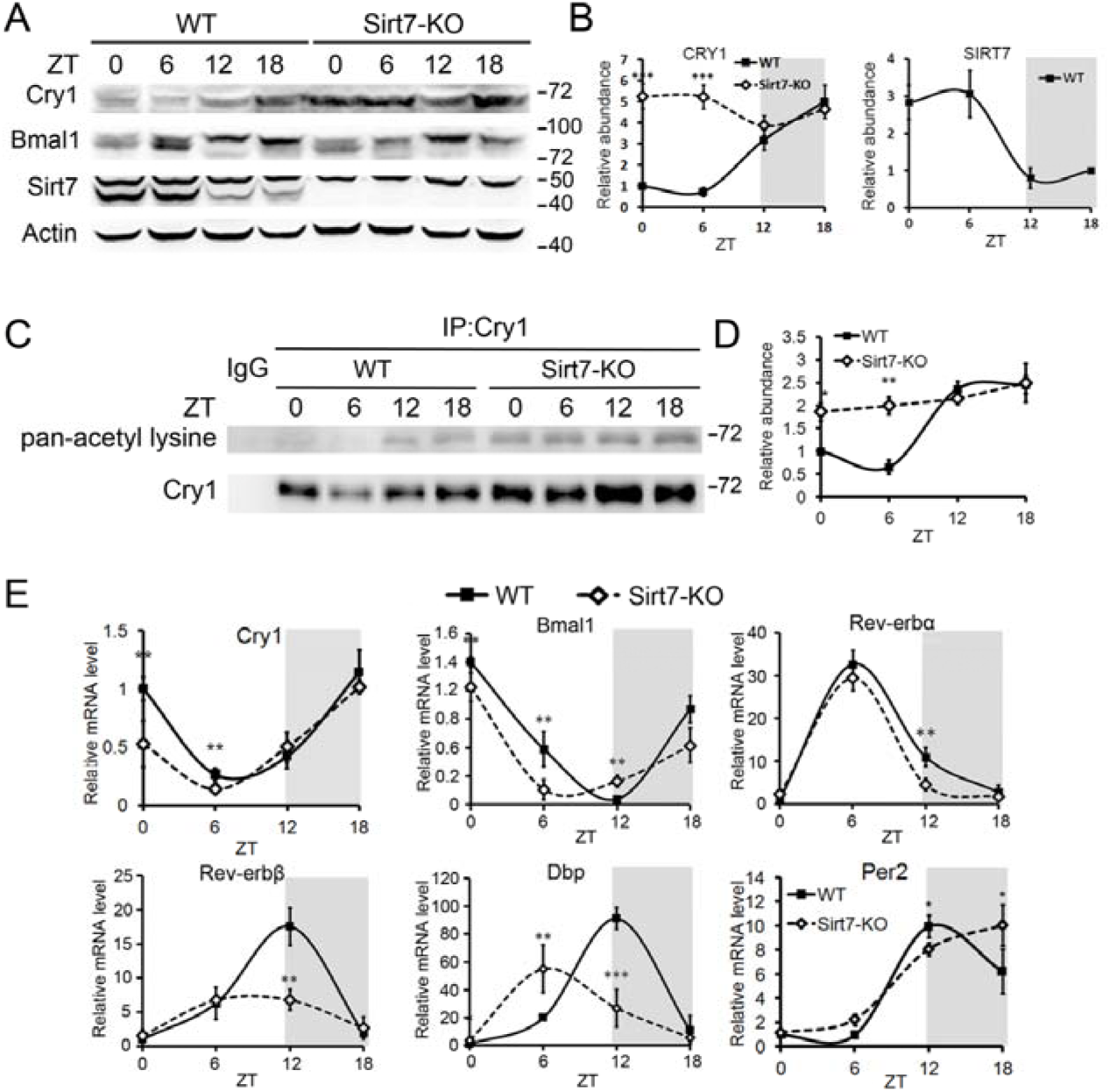
Sirt7 regulates circadian clocks in mouse livers. (***A***) Core clock protein oscillations in WT and *Sirt7*-/- liver tissues (n = 3 mice/genotype/time point). Representative blots from three independent experiments are shown. (***B***) Quantification of Cry1 and Sirt7 levels by Image J^®^ in (**A**). Data represent the means ± SEM. \*\*\**P* < 0.001. (***C***) Acetylation of Cry1 in WT (left) and *Sirt7*-/- (right) mouse livers (n = 3 mice/genotype/time point). Liver extracts from WT and *Sirt7*-/- mice were immunoprecipitated with Cry1 antibody. (***D***) Quantification of acetylation of Cry1 of three independent Western blots in (**C**). Data represent the means ± SEM. (*E*) Real-time PCR analysis of circadian clock genes in WT and *Sirt7*-/- liver tissues isolated from *Sirt7*-/- mice generated by CRISPR/Cas9-mediated genome editing (n = 4 mice/genotype/time point). Data represent the means ± SEM of three independent experiments. \**P* < 0.05, \*\**P* < 0.01, \*\*\**P* < 0.001.

Next, we examined mRNA levels of core clock genes in *Sirt7*-/- livers. As shown, *Sirt7* deficiency slightly attenuated mRNA levels of *Bmal1*, *Rev*-*Erbβ* and *Dbp*, but not *Per2* and *Rev*-*Erbα* (Fig. 6E). Though the amplitude of the circadian oscillation of *Cry1* was merely changed, its mRNA level was significantly reduced specifically in light phase in *Sirt7*-/- livers. Of particular interest, the circadian phase of *Bmal1*, *Cry1*, *Rev*-*Erbα* and *Dbp* was substantially advanced, whereas that of *Per2* was delayed in *Sirt7*-/- livers compared to WT. No phase shift of *Bmal1*, *Dbp* or *Per2* was observed in the hypothalamus of *Sirt7* KO mice (Supplemental Fig. S10). Consistent with that of mRNA, the circadian phase of Bmal1 protein was advanced in *Sirt7* KO livers. These suggest that Sirt7 regulates circadian phase of hepatic clocks.

Circadian phase of peripheral clock is determined by systemic cues from the central pacemaker and feeding activity. The SCN-orientated synchronizing signal counteracts feeding-induced phase entrainment of peripheral clock (Saini et al., 2013). Given the critical role of Sirt7 in SCN-driven synchronization of hepatic clock and the advanced circadian phase of *Sirt7*-/- livers, we reasoned that Sirt7-mediated synchronization might counteract phase entrainment of hepatic clock induced by feeding. To test this hypothesis, *Sirt7* KO and WT mice were fed *ad libitum* for three weeks, and then subjected to restricted feeding regimen with food available only during daytime. Mice were sacrificed every 4 h after the inversion of feeding regimen on the second and fourth day. The kinetics of clock gene phase shifting were determined by analyzing mRNA levels using realtime quantitative PCR (Fig. 7). After 2 days’ food shifting, circadian phase of *Bmal1*, *Cry1*, *Rev*-*erbα*, *Rev*-*erbβ* and *Dbp* were slightly changed in WT. Intriguingly, these genes adapted more rapidly in *Sirt7*-/- mice. For instance, *Bmal1*, *Cry1* and *Rev*-*erbα* levels showed up to 8 h phase advance in *Sirt7*-/- mice compared to WT, whereas the difference was less than 4 h in mice fed *ad libitum*. Although two peak expression of *Per2* was observed at ZT4 and ZT16 in WT mice, a new phase of *Per2* already appeared in *Sirt7* KO mice peaking at ZT4. After 4 days’ food shifting, the circadian phases of examined clock genes were completely reversed, and little difference was observed between *Sirt7* KO and WT mice. Collectively, the data suggest that Sirt7 counteracts the synchronization of hepatic clock to the feeding cues.

**Figure 7.**
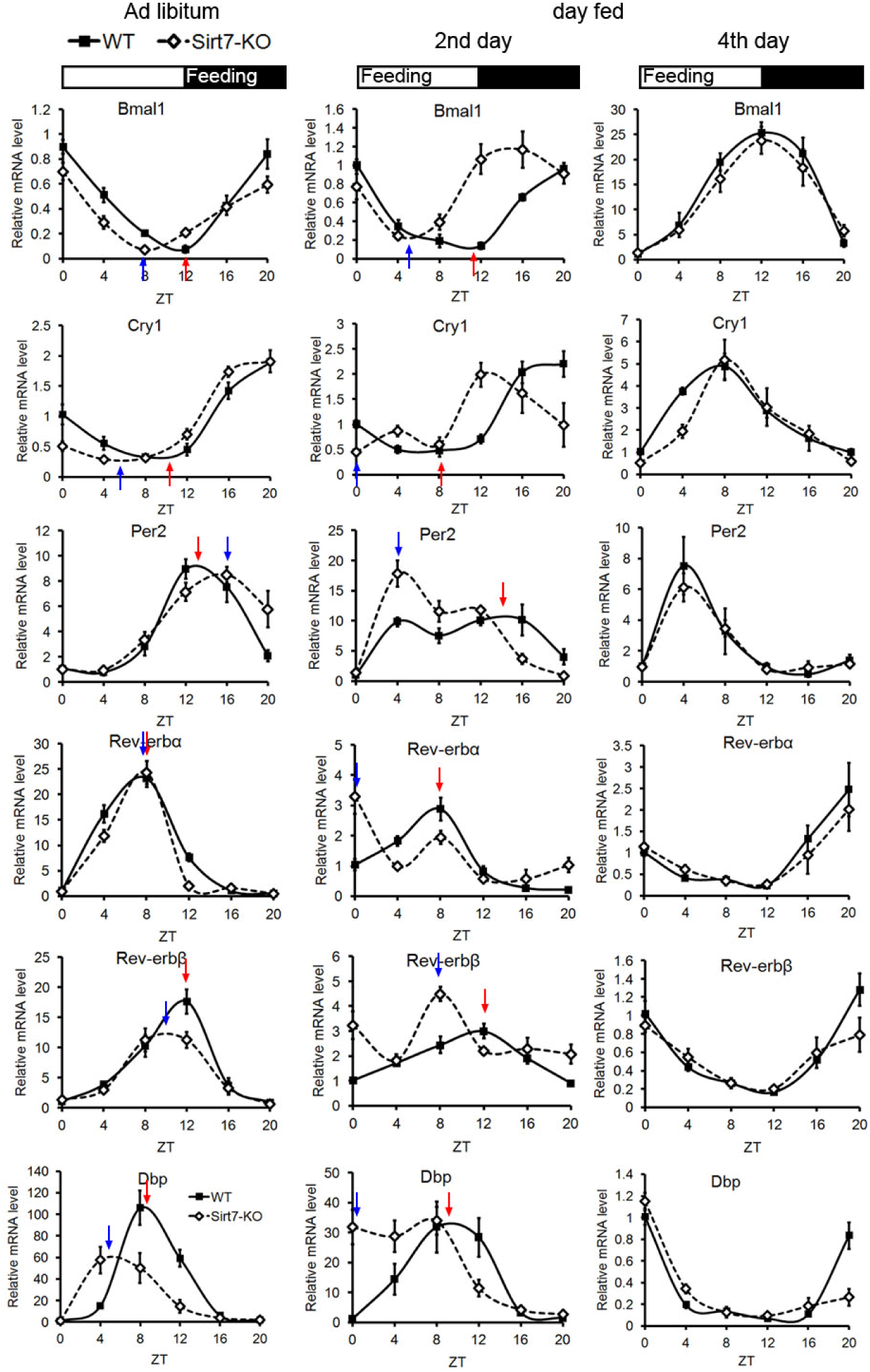
Sirt7 counteracts phase-shifting of hepatic clock induced by restricted feeding. Real-time PCR analysis of mRNA levels of core clock genes in WT and *Sirt7* KO mice fed *ad libitum* or exclusively during daytime (n = 3 mice/genotype/time point). Arrows indicate the maximal or minimum expression of the respective genes in an examined circadian cycle.

Altogether, we showed that BT cycles induce rhythmic transcription of Hsp70, which interacts with and promotes ubiquitination and proteasomal degradation of Sirt7. Sirt7 specifically deacetylates and promotes ubiquitination and degradation of Cry1, thus integrating hepatic clock to the central pacemaker. These results highlight Sirt7 as an early element responsive to light, thus transmitting timing information from the SCN to the periphery via the body temperature/Hsp70-Sirt7-Cry1 axis (Fig. 8).

**Figure 8.**
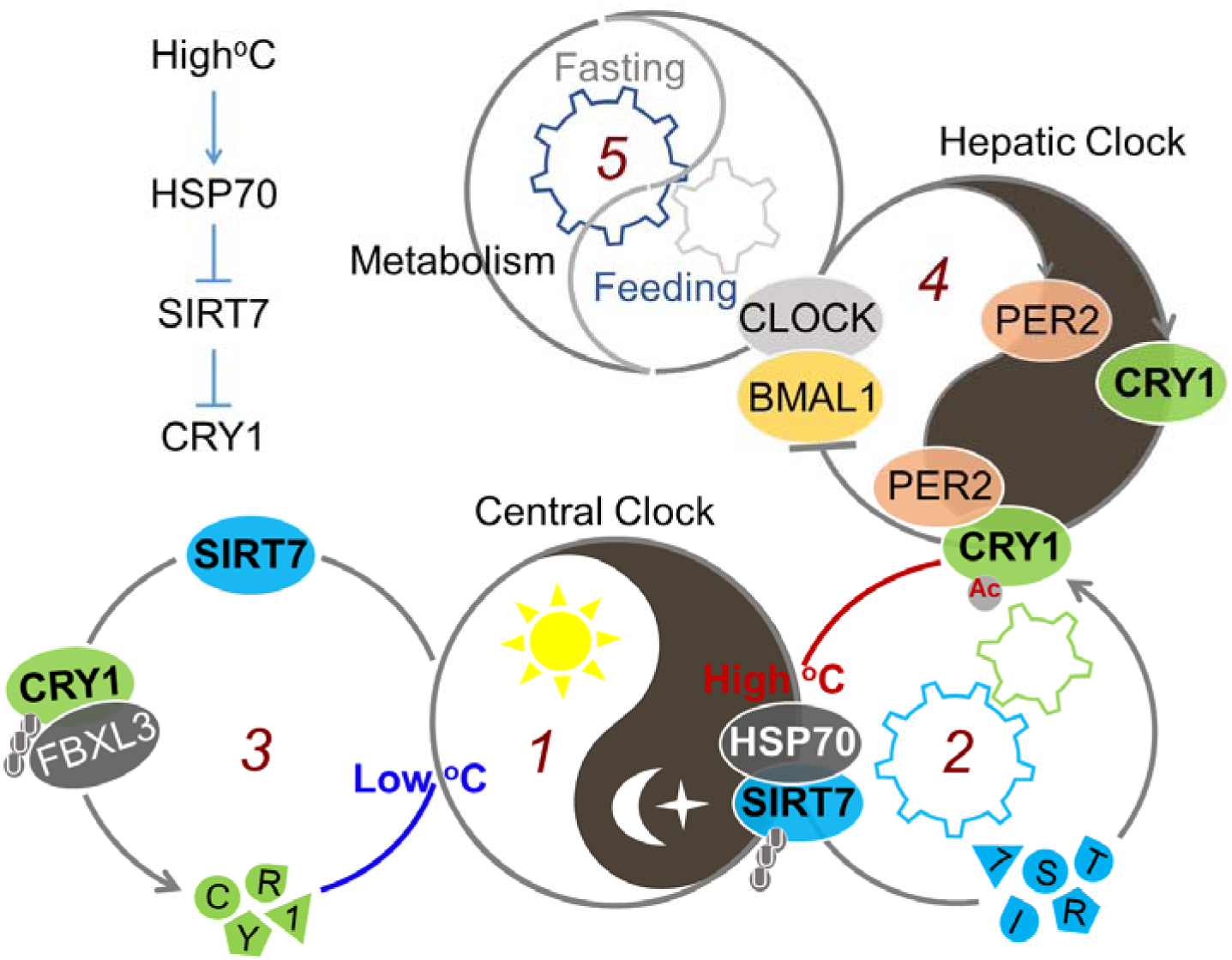
The BT/Hsp70-Sirt7-Cry1 axis ensures phase coherence between the central and peripheral clocks. A light-entrained BT cycle induces rhythmic expression of Hsp70 in the liver (Cycle 1), which interacts with Sirt7 and promotes its proteasomal degradation (Cycle 2), thus synchronizing hepatic clock. When abundant, Sirt7 deacetylates Cry1 to promote FBXL3-mediated ubiquitination and subsequent degradation (Cycle 3), thus coupling the central pacemaker to hepatic clock (Cycle 4). The BT/Hsp70-Sirt7-Cry1 axis-mediated synchronization of hepatic clock is independent of FF cycle (Cycle 5), which rather counteracts FF-entrained circadian phase in the liver.

## Discussion

The light-entrained central pacemaker in the SCN synchronizes peripheral clocks to establish a coherent circadian phase. Disruption of phase coherence leads to metabolic disturbances, diseases and even aging (Arble et al., 2009; Chang and Guarente, 2013; Turek et al., 2005). However, the molecular mechanisms, by which peripheral clocks are synchronized to the SCN, are far from clear. Here, we showed that Sirt7 in the liver is stabilized by light exposure but independent of core clock elements *Bmal1* and *Per2*, implicating Sirt7 as an early element responsive to light signaling and transmitting the timing information to mouse liver. Intriguingly, *Sirt7* is hardly detected in the hypothalamus (Supplemental Fig. S1C); depletion of *Sirt7* shifts circadian phase of the liver but not the hypothalamus, therefore desynchronizing the peripheral oscillators from the central pacemaker. Previous studies suggest that *Bmal1* and *Per2* are separately controlled by the SCN and food-derived resetting cues (Asher et al., 2010; Saini et al., 2013). In line with this notion, Sirt7 was reduced at ZT16 by light exposure in fasted mice, supporting an essential role for the LD cycle rather than food availability in driving rhythmic expression of Sirt7.

BT is a common resetting cue utilized by the SCN to entrain peripheral clocks (Brown et al., 2002; Buhr et al., 2010; Kornmann et al., 2007; Saini et al., 2012). Hsf1 is essential in resetting peripheral clocks via BT oscillation, as the transcription of *Hsps* is driven by Hsf1 in a circadian manner (Kornmann et al., 2007; Saini et al., 2012). However, at molecular levels, the connection between BT and local clock components are still largely unknown. Our results indicate that Hsp70, a key molecular chaperone, interacts with Sirt7, triggering poly-ubiquitination and proteasomal degradation, and Sirt7 thus integrates BT to hepatic clocks. Of note, a low core BT extends lifespan in mice, correlating with the anti-aging effect of calorie restriction (Conti et al., 2006; Heilbronn et al., 2006; Lane et al., 1996; Roth et al., 2002). Our data showed that low BT stabilizes Sirt7, therefore whether Sirt7 is involved in the BT-aging/longevity axis is worth further investigation.

Mammalian Crys share conserved photolyase-homology region (PHR) at N-terminus, but divergent carboxyl-terminal tail domain (CTD) (Czarna et al., 2013; Khan et al., 2012; Tamanini et al., 2007). Though generally accepted that CTD of Crys is crucial for the functional diversity (Chaves et al., 2006; Tamanini et al., 2007), the molecular mechanisms are not well understood. Here, we found that Sirt7 deacetylates Cry1 at K565/579, which are 2 of 10 Ks of Cry1 targeted for Fbxl3-mediated ubiquitination and subsequent degradation (Godinho et al., 2007; Hirano et al., 2013; Yoo et al., 2013). Constitutive high protein level of Cry1 in *Sirt7* null cells suppresses its own transcription, specifically at the light phase of a circadian cycle. Moreover, our results demonstrate that Cry1 but not Cry2, is deacetylated by Sirt7, as K565/579 residues are only present in the CTD of Cry1. Thus, our data shed new light on the function importance of CTD for Cry1 stability mediated by Sirt7 deacetylase. Although at this stage we cannot conclude that the altered kinetics of circadian phase shifting in *Sirt7* null mice are solely attributable to the elevated Cry1 level, similar to *Sirt7* deficiency, delayed circadian phase of *Per2* was observed in *Fbxl3*-/- livers (Shi et al., 2013; Siepka et al., 2007), supporting the notion that the disrupted circadian phenotype of *Sirt7*-/- mice is, at least partially, attributable to dysregulated Cry1. Notably, protein stability of Cry1 is also regulated by AMPK-mediated phosphorylation (Lamia et al., 2009). Our data suggest that Sirt7 and AMPK regulate Cry1 stability via independent pathways, involving ubiquitination at different residues.

It is hypothesized that SCN-derived signal counteracts food-induced phase-shifting of peripheral clocks (Le Minh et al., 2001; Saini et al., 2013). Our data support this hypothesis as the circadian phase-shifting response to feeding adapts more rapidly in *Sirt7* KO livers. Importantly, we reveal a novel molecular network, wherein Sirt7, as a central component, couples BT cues to hepatic oscillators. However, the physiological consequence of the altered kinetics of circadian phase shifting in *Sirt7*-/- mice is still unknown. It is previously reported that high-fat diet alters the feeding regimen, i.e. increased food intake during daytime; day-time feeding regimen results in metabolic syndrome in mice (Kohsaka et al., 2007; Mukherji et al., 2015a; Mukherji et al., 2015b). In contrast, restricted feeding during night-time rather protected mice from high-fat diet induced metabolic diseases (Hatori et al., 2012). Thus, one can speculate that the metabolic disturbances might be attributable to phase incoherence between the SCN and peripheral clocks. *Sirt7*-/- mice would be a useful model to elucidate metabolic pathways that are controlled by the SCN to counteract feeding cues. In conclusion, our data indicate that Sirt7 transmits light cues arriving at the SCN to peripheral via the BT/Hsp70-Sirt7-Cry1 axis.

## Experimental procedures

### Animals

*Sirt7* KO alleles were created by CRISPR/Cas9-mediated genome editing in C57BL/6 mice (transgenic animal services from Cyagen, Guangzhou, China). Briefly, the Cas9 mRNA and Sirt7-gRNA were generated by *in vitro* transcription and co-injected into fertilized eggs. Successful deletion was confirmed by PCR and DNA sequencing. *Sirt7* heterozygous males were backcrossed with C57BL/6 females for at least five generations to preclude off-targeted mutations. The sequence of the gRNA was as follow, 5’-CTTGGCCGAGAGCGAGGATC-3’. Mice were maintained on a 12 h LD cycle at 22°C with access to food and water *ad libitum*. Male mice aged 8-12 weeks were used in all experiments. For temperature challenges, mice were housed at 22°C for at least 4 weeks and then acutely shifted to 32°C (ZT0 to ZT4 or ZT12 to ZT16) or 4°C (ZT14 to ZT16) environment. Rectal temperatures were measured using a Thermocouple Meter (Landwind medical Industry Co., Ltd). All animals were housed and handled in accordance with protocols approved by the Committee on the Use of Live Animals in Teaching and Research of Shenzhen University.

### Cell culture

HEK293T cells were cultured in DMEM (Life Technologies), supplemented with 10% fetal bovine serum (FBS), 100 units/ml penicillin and 100 mg/ml streptomycin, and maintained in a humidified incubator at 37°C with 5% CO_2_. Primary MEFs were prepared from E13.5 embryos as described (Liu et al., 2005), grown in DMEM supplemented with 15% FBS, 2 mM glutamine (Life Technologies, US), 8 mM nonessential amino acids (Life Technologies, US), 1 mM Sodium Pyruvate (Life Technologies, US) and 10 mM HEPES (pH 7.0). Cells were transfected with indicated constructs using Lipofectamine^®^3000 reagent (Invitrogen, US), according to the manufacturer’s protocols. Cycloheximide (CHX, 50 μg/ml, Sigma-Aldrich), MG132 (10 μ, 6 h, Sigma-Aldrich), nicotinamide (NAM, 10 mM, 6 h, Sigma-Aldrich) were added to the cultures as indicated.

### Plasmids

Construct FLAG-Sirt7, FLAG-Cry1 HA-Sirt7 and HA-Cry1 were constructed as follow: the full length coding sequence of human Cry1 and Sirt7 was amplified from cDNA of HEK293 cells by RT-PCR; the PCR products were cloned between the *Xho*I and *Xba*Ior *Bam*HI and *Xho*Isites in pcDNA3.1 vector. FLAG-Sirt7 H187Y was purchased from Addgene (53151). FLAG-His-Hsp70, FLAG-His-CRY2 and FLAG-His-FBXL3 were purchased from ViGene Biosciences Inc. Various KR mutations were introduced by PCR-based site-directed mutagenesis. Myc-Ub construct was a gift from Dr. Z.J. Zhou (School of Biomedical Sciences, LKS Faculty of Medicine, the University of Hong Kong).

### CRISPR/Cas9 mediated gene deletion

CRISPR/Cas9-mediated gene deletion was conducted as previously described (Ran et al. 2013). In brief, guide RNA targeting Sirt7 was subcloned into PX459 vector (Addgene#48139). HEK293T cells were transfected with 1 μg PX459-gSirt7. After 48 h, clones were selected with 1 μg/ml puromycin (Invitrogen), then expanded for further analysis. The mutations were confirmed by PCR, DNA sequencing and immunoblotting. To avoid non-specific off-target effect, two independent gRNAs were applied, the sequences are as follow,

gSirt7-1-F: 5’-CACCGCCGCCGCTTTGCGCTCGGAG-3’,

gSirt7-1-R: 5’-AAACCTCCGAGCGCAAAGCGGCGGC-3’;

gSirt7-2-F: 5’-CACCGTGTGTAGACGACCAAGTATT-3’,

gSirt7-2-R: 5’-AAACAATACTTGGTCGTCTACACAC-3’.

### Antibodies, Immunoprecipitation and Western blotting

Anti-FLAG and Anti-HA antibodies were from Sigma. Antibodies for Immunoblotting include anti-SIRT1 (#8469, Cell Signaling Technology), anti-SIRT2 (ab67299, Abcam), anti-SIRT3 (#5490, Cell Signaling Technology), anti-SIRT4 (Sigma), anti-SIRT5 (ab108968, Abcam), anti-SIRT6 (NB100-2252, Novus), anti-Sirt7 (sc-135055, Santa Cruz Biotechnology), anti-BMAL1 (ab93806, Abcam), anti-Cry1 (ab3518, Abcam), anti-PER2 (sc-25363, Santa Cruz Biotechnology), anti-Hsp70 (#4872, Cell Signaling Technology), anti-Ubiquitin (#3936, Cell Signaling Technology), anti-α-TUBLIN, anti-GST (2624, Cell Signaling Technology), anti-His and pan acetyl lysine antibody (PTM-105, PTM-Biolab). Antibodies used for immunoprecipitation are anti-FLAG M2 Affinity Gel (Sigma), monoclonal anti-HA Agarose (Sigma), anti-Sirt7 (sc-365344, Santa Cruz Biotechnology) and anti-Cry1 (ab3518, Abcam).

Whole-cell extracts were prepared in IP lysis buffer (20 mM Tris-HCl, pH 8.0, 250 mM NaCl, 0.2% NP-40, 10% glycerol, 2 mM EDTA, 1 mM PMSF, and Protease inhibitor cocktail). For immunoprecipitation, whole-cell lysates were mixed with 2 μg primary antibody as indicated, or control IgG and precipitated by Protein A/G agarose beads (Pierce). To evaluate acetylation of Cry1, whole-cell extracts were prepared in IP lysis buffer supplemented with 10 mM sodium butyrate (NaB) and 10 mM NAM. Equal amounts of cell lysates or immunoprecipitated protein samples were separated by SDS-PAGE and transferred to a PVDF membrane (Millipore). Primary antibodies were incubated at 4°C overnight. After incubation with secondary antibodies conjugated to HRP (Jackson lab), the signal was visualized using an enhanced ECL chemiluminescence detection system (Sage Creation Science).

### RNA extraction and Real-time PCR

Total RNA were extracted with TRIzol^®^ reagent (Invitrogen). Complementary DNA was synthesized from 2 μg RNA using Primescript^®^ RT Master Kit (Takara, Japan) according to the manufacturer’s instructions. Quantitative RT-PCR was performed on a BIO-RAD CFX Connect^™^ Real-time PCR system with SYBR Ex Taq Premixes (Takara, Japan). The relative quantification Delta-delta Ct method was used for analysis. Mouse *36b4* and human *ACTIN* were used as internal controls to normalize all data. Primers are listed in Table S1

### GST-pull down assay

GST-pull down assay was performed using recombinant GST-Sirt7 and His-Cry1 purified from BL21 *E. coli*. GST or GST-Cry1 protein (2 μg) was immobilized on Glutathione-Sepharose 4B, and incubated with His-Cry1 in GST binding buffer (20 mM Tris-HCl [pH 7.4], 0.1 mM EDTA, and 150 mM NaCl, 0.2% NP-40, protease inhibitors cocktail). The beads were washed three times with GST wash buffer (20 mM Tris-HCl [pH 7.4], 0.1 mM EDTA, and 250 mM NaCl, 0.2% NP-40). Bead-bound Cry1 was analyzed by SDS-PAGE and western blotting. For purification of recombinant His-Cry1 from BL21 *E. coli*, 1 mM FAD (flavin adenine dinucleotide) were added in the lysis buffer (50 mM PBS [pH 7.4], 0.5 M NaCl, 1 mM PMSF, protease inhibitors cocktail).

### *In vitro* deacetylation assay

FLAG-Cry1 was over-expressed in HEK293T cells and immunoprecipitated by Anti-FLAG M2 Affinity Gel (Sigma). For deacetylation assay, purified FLAG-Cry1 was incubated with 1 μg GST-Sirt7 in deacetylation buffer (50 mM Tris-HCl [pH 8.0], 4 mM MgCl2, 0.2 mM DTT, 1 mM NAD^+^, protease inhibitors cocktail) for 30 min with constant agitation. The acetylation level of Cry1 was monitored by Western blotting using anti-acetyl lysine antibodies.

### Mass spectrometry

Gel lanes were cut from the gel and subjected to in-gel digestion with trypsin. Digested peptides were resuspended and analyzed by LC-MS with a QTRAP 6500 mass spectrometer (Applied Biosystems). Data analysis was performed using the Mascot search engine against IPI-HUMAN and NCBI databases for protein identification.

## Acknowledgements

This study was supported by grants from National Natural Science Foundation of China (81422016, 91439133, 81571374 to B.L., and 81601215 to Z.L.), National Key R&D Program of China (2017YFA0503900 and 2016YFC0904600), Guangdong Province (2014A030308011 to B.L.), Shenzhen (CXZZ20140903103747568 and JCYJ20160226191451487 to B.L., and Discipline Construction Funding of Shenzhen). The authors would like to thank Dr. Jessica Tamanini (Shenzhen University and ETediting) for editing the manuscript prior to submission.

